# Eliminating redundancy among protein sequences using submodular optimization

**DOI:** 10.1101/051201

**Authors:** Maxwell W. Libbrecht, Jeffrey A. Bilmes, William Stafford Noble

## Abstract

**Motivation:** Submodular optimization, a discrete analogue to continuous convex optimization, has been used with great success in many fields but is not yet widely used in biology. We apply submodular optimization to the problem of removing redundancy in protein sequence data sets. This is a common step in many bioinformatics and structural biology workflows, including creation of non-redundant training sets for sequence and structural models as well as selection of “operational taxonomic units” from metagenomics data.

**Results:** We demonstrate that the submodular optimization approach results in representative protein sequence subsets with greater structural diversity than sets chosen by existing methods. In particular, we compare to a widely used, heuristic algorithm implemented in software tools such as CD-HIT, as well to as a variety of standard clustering methods, using as a gold standard the SCOPe library of protein domain structures. In this setting, submodular optimization consistently yields protein sequence subsets that include more SCOPe domain families than sets of the same size selected by competing approaches. We also show how the optimization framework allows us to design a mixture objective function that performs well for both large and small representative sets. The framework we describe is theoretically optimal under some assumptions, and it is flexible and intuitive because it applies generic methods to optimize one of a variety of objective functions. This application serves as a model for how submodular optimization can be applied to other discrete problems in biology.

**Availability:** Source code is available at https://github.com/mlibbrecht/submodular_sequence_repset.

## 1 Introduction

Redundancy is ubiquitous in biological sequence data sets, and this redundancy often impedes analysis. For example, because some proteins, such as those involved in disease or industrial applications, are studied by many more researchers than proteins with less important or unknown function, protein sequence databases contain hundreds or thousands of slight variants of well-studied proteins and only one or a few copies of the sequences of less-studied proteins. Removing this redundancy has two benefits. First, doing so removes computational and statistical issues introduced by this redundancy. Second, members of a non-redundant set of sequences can be interpreted as representatives of the whole set, such as using sequences to represent species or “operational taxonomic units” in a metagenomics study [Human Microbiome Project Consortium, 2012].

Redundancy can be handled by selecting a *representative subset* of the original data set. This representative subset is then used in downstream analysis. However, the task of choosing this subset can be computationally challenging because finding the best representative subset out of *N* items requires, in principle, considering all *2*^*N*^ possible subsets.

In many fields, representative subsets are selected using some form of *submodular optimization*. The class of submodular functions is analogous to the class of convex functions. Submodular functions, however, are defined over subsets of a given set of data items, whereas convex functions are defined over real-valued vectors. Submodular functions are those that satisfy the property of diminishing returns. If we think of a function *f* as measuring the quality of a given subset, then the submodular property means that the incremental “value” of an item *v* decreases as the context in which *v* is considered grows. Formally, for any two subsets *A* and *B* of the set *V*, where *A* ⊆ *B* ⊂ *V*, and for any *v* ∈ *V \ B*, *f* is submodular if and only if it satisfies *f*(*A* ⋃{*v*}) – *f*(*A*) ≥ *f*(*B* ⋃ {*v*}) – *f*(*B*). In many applications, it is common to search for a subset of maximal quality, as measured by a function *f*. The resulting optimization problem is hopelessly difficult for an arbitrary set function, but when the set function is submodular, the quality can be maximized within a constant factor of optimal in low-order polynomial time [Fisher et al., 1978, Nemhauser et al., 1978, Minoux, 1978] (Methods). Moreover, these optimization algorithms are provably the best achievable in polynomial time under some assumptions (Methods). For these reasons, submodular optimization has a long history in economics [Vives, 2001, Carter, 2001], game theory [Topkis, 1998, Shapley, 1971], combinatorial optimization [Edmonds, 1970, Lovász, 1983, Schrijver, 2004], electrical networks [Narayanan, 1997], and operations research [Cornunéjols et al., 1990]. Furthermore, submodular optimization has recently been used with great success for selecting representative subsets of text documents [Lin et al., 2007, Lin and Bilmes, 2011, 2012], recorded speech [Liu et al., 2013, Wei et al., 2013, 2014], machine translation [Kirchhoff and Bilmes, 2014], and images [Tschiatschek et al., 2014]. However, submodular optimization is not yet widely used in biology.

To illustrate how submodular optimization can be applied to biological data sets, we focus on the problem of choosing representative subsets from a database of protein sequences. Many methods have been developed for this problem [Hobohm et al., 1992, Holm and Sander, 1998, Li et al., 2001, Edgar, 2010, Parsons et al., 1992, Altschul et al., 1997, Enright and Ouzounis, 2000, Rice et al., 2000], including the popular CD-HIT [Weizhong and Adam, 2006] and PISCES [Wang and Dunbrack, Jr., 2003] algorithms. These sequence selection methods are very widely used— for example, the CD-HIT papers have been cited a total of >3,000 times (Google Scholar)—and are a standard preprocessing step applied to data sets of protein sequences, cDNA sequences and microbial DNA. All of these existing methods are based on the following simple algorithm: start with an empty set; order the sequences by length; then for each sequence, add the sequence to the set if no sequence currently in the set is more similar to it than some threshold (usually 90% sequence identity). Despite the ubiquity of this algorithm, it is simply a heuristic, apparently without any theoretical or empirical justification for its use over any other.

We propose a principled framework for representative protein sequence subset selection using submodular optimization. This approach involves defining a submodular objective function that quantifies the desirable properties of a given subset of sequences, and then applying a submodular optimization algorithm to choose a representative subset that maximizes this function. For illustrative purposes, we applied this optimization framework to protein domain sequences drawn from a single class (“Membrane and cell surface proteins and peptides”) within the Structural Classification of Proteins—extended (SCOPe) database [Murzin et al., 1995]. The availability of protein structures provide an orthogonal gold standard for judging the quality of a given representative subset. The results (Figure 1) suggest that the submodular approach does a good job of selecting protein sequences with diverse structural properties. In particular, among the nine folds that comprise our selected class, the submodular method successively chooses one exemplar sequence from fold before selecting two sequences from the same fold. The only exception is that the method chooses two “transmembrane beta-barrel” protein domains before selecting the first “single transmembrane helix” domain; however, as illustrated in Figure 1, this choice is not particularly surprising, given that the “transmembrane beta-barrel” fold is comprised of two superfamilies with quite different structural properties.

**Figure 1:**
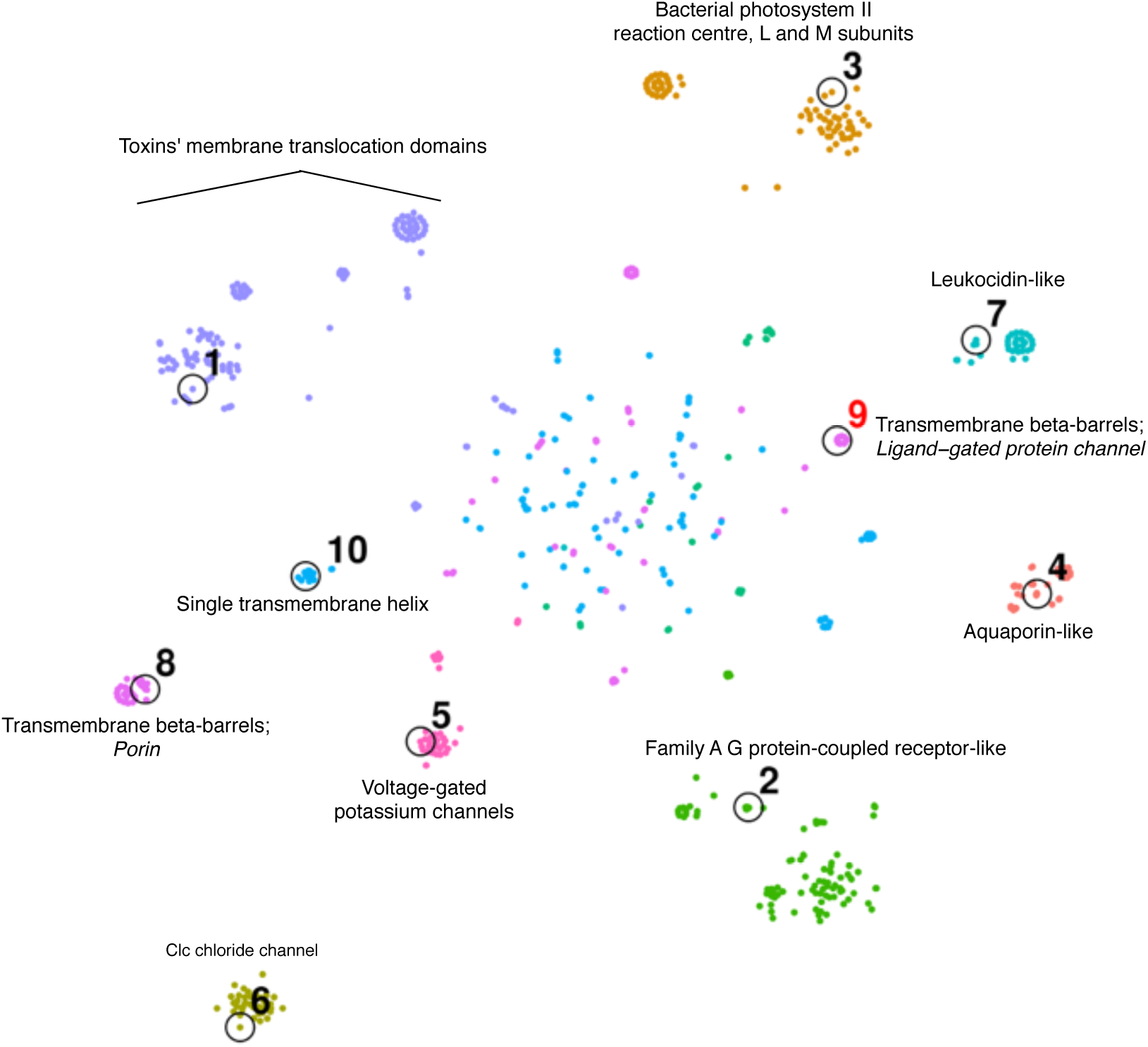
Protein sequence representative set selection. Points represent protein sequences, projected into 2D based on their pairwise similarities according to BLAST using t-SNE [Van der Maaten and Hinton, 2008]. All sequences are depicted from the SCOPe Class “Membrane and cell surface proteins and peptides.” Color with text labels indicates SCOPe fold, and italicized text indicates SCOPe family within a given fold. Black circles indicate the top ten representative sequences chosen by the greedy algorithm on a mixture of facility-location (Rankprop sim; 0.5 weight) and sum-redundancy (percent ID sim; 0.5 weight). Numbers labeling these circles indicate choice order. Choice #9 is colored red because it is the second chosen sequence within the same fold. To produce the 2D embedding, we first computed *a n × n* pairwise distance matrix (for a set of *n* sequences) with the formula: distance(*i, j*) = 1 – percent_id(*i,j*)/100. We used multidimensional scaling (MDS) to project this distance matrix to a 30 × *n* feature matrix. This MDS preprocessing step is similar to the standard preprocessing step used by tSNE that applies principle component analysis (PCA) to project a high-dimensional feature matrix to a 30 × *n* feature matrix [Van der Maaten and Hinton, 2008]. We used tSNE to project this 30 × *n* matrix into a 2 × *n* matrix, which is plotted. The diffuse points in the center are a common feature of tSNE plots, and are a consequence of the algorithm imperfectly recapitulating the *n*-dimensional similarity matrix in two dimensions.

In the remainder of this paper, we expand upon this example, demonstrating that submodular optimization method does a better job of selecting representative protein sequence sets than competing methods, when evaluated relative to protein structure gold standards. We also demonstrate how optimization-based method development offers practical advantages over an approach in which the algorithm and the property it tries to optimize (its objective function) are inextricably intertwined. In particular, we demonstrate how a hybrid optimization function allows us to design a method that excels at selecting either large or small representative sets.

Note that the task of representative subset selection is distinct from that of clustering. Clustering focuses on sets of similar sequences, whereas representative subset selection focuses on the individual sequences chosen to represent a larger set. A large number of methods have been proposed for protein sequence clustering, including those based on Markov clustering [Enright et al., 2002], connected component detection [Enright and Ouzounis, 2000], generative modeling [Yang and Wang, 2003], minimal spanning trees [Guan and Du, 1998], and spectral clustering [Paccanaro et al., 2006]. Applying these methods to representative subset selection requires specifying, in addition, a procedure for selecting an exemplar sequence from each cluster. In practice, clustering methods are rarely used for reducing redundancy in protein sequence sets. Nonetheless, we empirically demonstrate below that our submodular selection approach outperforms several commonly used clustering algorithms.

## 2 Methods

### 2.1 Submodularity

The property of *submodularity* is important for the optimization of the set functions defined below. A *submodular* function [Fujishige, 2005] is defined as follows: given a finite set *V* = {1, 2,…, *n*}, a discrete set function *f*: 2^*V*^ → ℝ is submodular if and only if:

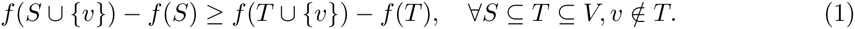

In other words, the incremental gain of adding item *s* to the set decreases when the set to which *s* is added to grows from *S* to *T*. Intuitively, most reasonable measures of the quality or “informativeness” of a set of items are submodular because the gain from adding a given item is reduced when the set already contains other items similar to it.

Two other properties of set functions are relevant to their optimization. First, a set function *f* is defined as *monotone non-decreasing* if

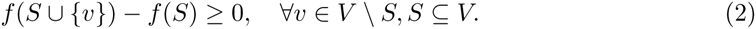

Second, we say that *f* is *normalized* if *f*(*ø*) = 0.

### 2.2 Similarity functions

The measures of subset quality described below are based on pairwise similarities of sequences. Such similarity measures are real-valued functions *s*(*u,v*) defined on pairs of sequences. We considered two measures to quantify the similarity between a pair sequences: (1) the fraction of residues that match between the two sequences (fraction identical), and (2) the similarity measure used by the Rankprop method, defined as exp(—E-value/100) [Weston et al., 2004]. Both measures have been used successfully in previous work on protein sequences, and can be calculated efficiently using an index-based method like BLAST [Altschul et al., 1990]. In both cases, we define the similarity between all pairs of sequences not reported by BLAST as 0. Note that, due to differing sequence lengths, both measures are not symmetric.

### 2.3 Subset quality measures

Let *V* be the full set of sequences and let *s*(*u,v*) be a measure of similarity between a pair of sequences *u,v* ∈ *V*. A subset quality function is defined over subsets *f*(*A*), where *A* ⊆ *V*. We consider several such subset quality functions, each of which measures some property of a set of sequences. In each case, a higher value of the function is desired.

First, we define the *facility-location* function as

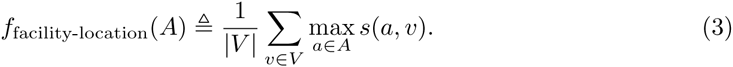

Intuitively, this function takes a high value when every sequence in *V* has at least one similar representative in *A*. This function is also the objective function of the *k*-medoids clustering method. The facility-location function is submodular, normalized and monotone non-decreasing.

Second, we define the *sum-redundancy* function as

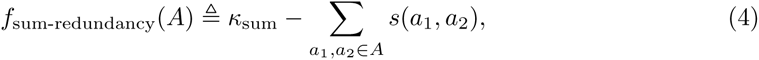

where *k*_sum_ = Σ_*a*_,_*b∈V*_*s*(*a, b*). Intuitively, for sets *A* of a given size |*A*|, this function takes a very small value when *A* includes many pairs of similar sequences, and it takes a large value when all the sequences in *A* are very dissimilar from one another. In other words, the sum-redundancy function penalizes the redundancy in *A*. Optimizing the sum-redundancy function with the monotone greedy algorithm (defined below) is identical to the commonly-used representative subset heuristic of iteratively choosing the example that is most dissimilar from all previously-chosen examples. The sum-redundancy is submodular and normalized, but not monotone non-decreasing (in fact, it is monotone non-increasing).

Third, we define the *max-redundancy* function as

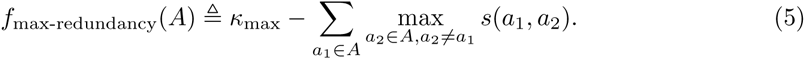

where *k*_max_ = Σ_*a*_1_∈*V*_ max_*a*_2_∈_*V, a*_2_≠*a*_1_ *s*(*a*_1_,*a*_2_). The max-redundancy function is very similar to sum-redundancy, except that it penalizes only the most similar neighbor for each sequence in *A* instead of all neighbors. The max-redundancy is submodular and normalized, but not monotone non-decreasing and (like above) is monotone non-increasing.

### 2.4 Optimization algorithms

We are interested in choosing a subset of sequences *A* that maximizes a given measure of quality of the subset. Because all of the set functions defined above are either monotone increasing or monotone decreasing in the size of the set, optimizing the function over all subsets would result in either the empty set or the full set, depending on the function. Therefore, we additionally control the size of the returned set using one of two strategies. In the first setting, we constrain the size of the returned set to a specific size *k* and solve the optimization problem

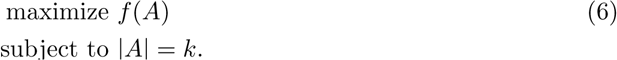

In the second setting, we add a term to the objective that is proportional to the size of the subset, scaled by a hyperparameter λ, and we optimize the unconstrained function

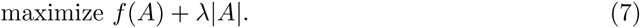

We use the constrained strategy when using the monotone greedy algorithm (defined below) because that algorithm supports such constraints. The bidirectional greedy algorithm (defined below) works only for unconstrained problems, so we use the second strategy for that algorithm, varying λ to indirectly control the resultant size. In practice, if a particular subset size (or some other feature of the set such as minimum objective value) is desired, this can be achieved using binary search.

Maximizing an arbitrary set function in general requires considering all 2^|*v*|^ possible subsets and is therefore impractical for large sets of sequences. In fact, even when this set function is submodular, maximizing it exactly is NP-hard. However, submodular functions can be efficiently approximated within a constant factor of optimal. In this work we apply the following two optimization algorithms.

The first algorithm is the *monotone greedy* algorithm (Supplementary Algorithm 2). At each iteration, the monotone greedy algorithm computes the difference in the objective function obtained by adding each sequence and adds the sequence with the largest difference. It turns out that when the objective function is submodular, monotone nondecreasing and normalized, this simple algorithm is guaranteed to produce a solution within a factor of 1 — 1/e ≈ 0.63 of optimal [Nemhauser et al., 1978]. This is the best approximation ratio achievable in polynomial time unless P=NP [Feige, 1998]. Furthermore, this algorithm can be sped up using the fact that the submodular property guarantees that, if adding a sequence *v* to *A* would result in a difference of *d* in the objective, then adding *v* to a larger set *AU B* will result in an objective difference *d*’ ≤ *d*. This property allows the algorithm to skip considering many sequences, and in practice often results in near-linear running time [Minoux, 1978].

The second algorithm is the *bidirectional greedy* algorithm (Supplementary Algorithm 3). This algorithm, which is randomized, maintains a growing set, which is initialized to the empty set, and a shrinking set, which is initialized to the full set. It considers each sequence *v* in turn, and either adds *v* to the growing set or removes it from the shrinking set randomly with probability proportional to the gain in the objective resulting from either change, respectively. At the end of the algorithm, the growing set and shrinking set are identical, and this set is returned. When the objective function is submodular and normalized (but not necessarily monotone nondecreasing), the mean of the objective values found over many randomized runs of this algorithm is guaranteed to be at least ½ of optimal [Buchbinder et al., 2012]. This is the best approximation ratio achievable in polynomial time (independent of whether P=NP) [Feige et al., 2011].

In the experiments below, we applied the monotone greedy algorithm to optimize the (monotone) facility-location function (1 — 1/e approximation guarantee). We applied the bidirectional greedy algorithm optimize the (nonmonotone) sum-redundancy and max-redundancy functions (1/2 approximation guarantee). The bidirectional greedy algorithm can be used on mixtures of these functions to maintain a 1/2 approximation guarantee. However, we achieved better results on the development data set using the greedy algorithm to optimize these mixtures (no theoretical guarantee; Supplementary Figure 8), so we used this strategy in our subsequent experiments.

Note that the 1/2 approximation guarantee can be maintained by running both algorithms and choosing the result with the better objective value. Moreover, although the greedy algorithm does not have a mathematical guarantee for non-monotone problems, we have found that for many problems it nonetheless performs better than the mathematically-guaranteed bidirectional greedy algorithm (results not shown). As future work, we are interested in investigating mathematical properties of these functions that may explain why the greedy algorithm is performing better than the current theory predicts.

### 2.5 Comparison to threshold algorithm (CD-HIT)

We call the algorithm used by most previous sequence subset selection methods, including CD-HIT and PISCES, the *threshold* algorithm (Supplementary Algorithm 1). This algorithm starts with an empty return set. It considers each sequence in decreasing order of sequence length, adding a given sequence to the return set if there is no sequence currently in the return set with similarity greater than some threshold *τ*. The value of *τ* controls the size of subset returned: a small *τ* results in a small set, while a large *τ* results in a large set.

In order to facilitate comparison, we implemented our own version of CD-HIT. The primary difference between our implementation and the CD-HIT source is that our implementation uses pre-computed BLAST alignments, while the CD-HIT source uses an on-the-fly index-based alignment method similar to BLAST. The main advantage of this on-the-fly method is that it does not require an expensive database construction preprocessing step. However, when comparing multiple methods on a single data set, it is much more efficient to perform a single alignment step. Using a single set of alignments also removes the possibility for performance differences due to slightly different alignment results. Note that the on-the-fly method could in principle be used in the context of either the CD-HIT or submodular optimization. Our implementation of CD-HIT and the CD-HIT source result in nearly-identical subsets (Supplementary Note 1).

### 2.6 Protein sequence data

To test these methods, we downloaded the full Structural Classification Of Proteins–extended (SCOPe) database version 15 from http://scop.berkeley.edu [Murzin et al., 1995]. SCOPe is a hierarchical structural classification database, consisting of four levels with increasing granularity: class, fold, superfamily, and family. In this work, we focus primarily on the family and superfamily levels because these represent a reasonable level of granularity for subset selection.

We also downloaded the ASTRAL database from http://scop.berkeley.edu [Brenner et al., 2000]. ASTRAL is a database of protein domain sequences, some of which are associated with SCOPe categories. We used only ATOM-derived sequence records, and we chose the subset of ASTRAL sequences that are associated with a SCOPe family, removing eight sequences that were labeled with multiple SCOPe families. This resulted in a set of 78,048 sequences.

We also downloaded the results from all-by-all BLAST on the ASTRAL sequences from http://scop.berkeley.edu. BLAST was run using the command line blastpgp-d DBNAME-i SEQ.fa-j 1-h 1e-2-v 5000-b 5000-a2-z 100000000-J T-O OUTPUT. This results in all pairwise relationships with an E-value less than 0.01, where the similarity of each pair is measured by an E-value and a fraction sequence identity.

In order to overfitting to SCOPe, we performed all development on just the 16,880 ASTRAL sequences with the SCOPe Class “All beta proteins.” Unless otherwise noted, the figures show performance on all of ASTRAL; we computed this performance only when preparing this manuscript.

## 3 Results

### 3.1 Submodular optimization effectively eliminates redundancy in sequence data sets

To evaluate how well submodular optimization can remove redundancy in protein sequence data sets, we applied it to ASTRAL sequences. We focus on the sum-redundancy function with percent identity objective function because this function is most comparable to CD-HIT and it performed best in our later evaluation (see below). As expected, submodular optimization produced representative subsets with less total redundancy (Figure 2A). This because submodular optimization is a theoretically-based algorithm that explicitly aims to minimize redundancy, while the threshold algorithm is a heuristic that lacks any guarantee. The improvement primarily comes from the fact that the threshold method chooses many pairs of sequences with identity just under the threshold, whereas optimizing sum-redundancy chooses very few sequences with any detectable identity at the cost of choosing a small number of closely-related pairs (Figure 2B). The small number of closely-related pairs that are accepted is likely due to the randomized nature of the algorithm.

**Figure 2:**
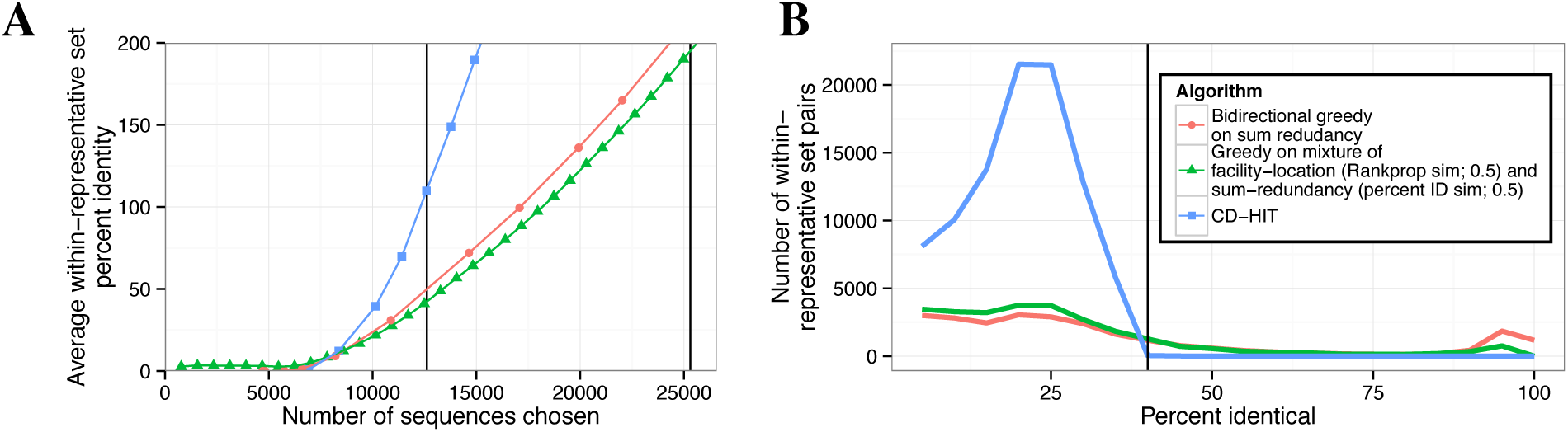
(A) Average redundancy in representative sets. The vertical axis is calculated for a representative set *A* as (1/|*A*|) Σ_*a*_1_,*a*_2_∈*A*_ *s*(*a*_1_,*a*_2_), where s(*a*_1_,*a*_2_) is percent identity. Vertical lines correspond to the sizes of subsets selected by CD-HIT using 40% and 90% sequence identity thresholds. (B) Histogram of pairwise similarities within representative sets. We produced subsets of size 12,614 sequences (the size reported by CD-HIT with a 40% identity threshold) for all methods by finding the smallest subset larger than the target size and randomly removing sequences to reach this size. Vertical axis indicates the number of pairs of sequences with a given percent identity within each size-12,614 representative set, in 5% bins. Vertical line indicates 40% identity threshold used by CD-HIT.

### 3.2 Representative subsets chosen by submodular optimization exhibit high structural diversity

In order to evaluate the quality of a subset of sequences independently of a specific objective function, we compared these subsets to the Structural Classification Of Proteins-extended (SCOPe) database. SCOPe contains 78,048 protein domain sequences, each assigned to one of 4,994 protein families. These assignments are based on experimentally determined protein structures, so they are independent from the sequence-based information used by the optimization methods. Intuitively, a good representative subset should include one sequence from each family, whereas a poor subset may include many sequences from the same family while leaving other families with no representatives. Therefore, we defined the quality of a subset as the number of SCOPe protein families with at least one sequence from that family in the subset.

Among the three submodular objective functions that we considered, we found that optimizing the sum-redundancy function produced the best results on this evaluation metric. Our evaluation considered 12 possible approaches: two submodular optimization algorithms on three different objective functions, each computed using two different measures of sequence similarity. To avoid overfitting to SCOPe, we performed all development and comparison of objective functions on a subset of the data (˜21%) and applied only the three best methods to the full data set. Optimizing sum-redundancy with the *bidirectional greedy algorithm* results in much better coverage of SCOPe families than the threshold heuristic method that is employed by methods such CD-HIT (Figure 3). For representative sets of size 12,614 (the size chosen by CD-HIT with threshold 40%), the set chosen by submodular optimization leaves 27% fewer SCOPe families uncovered (276 compared with 379) than the threshold method does. We reached a similar conclusion when performing the evaluation on SCOPe superfamilies and folds rather than families (Supplementary Figures 3-8).

**Figure 3:**
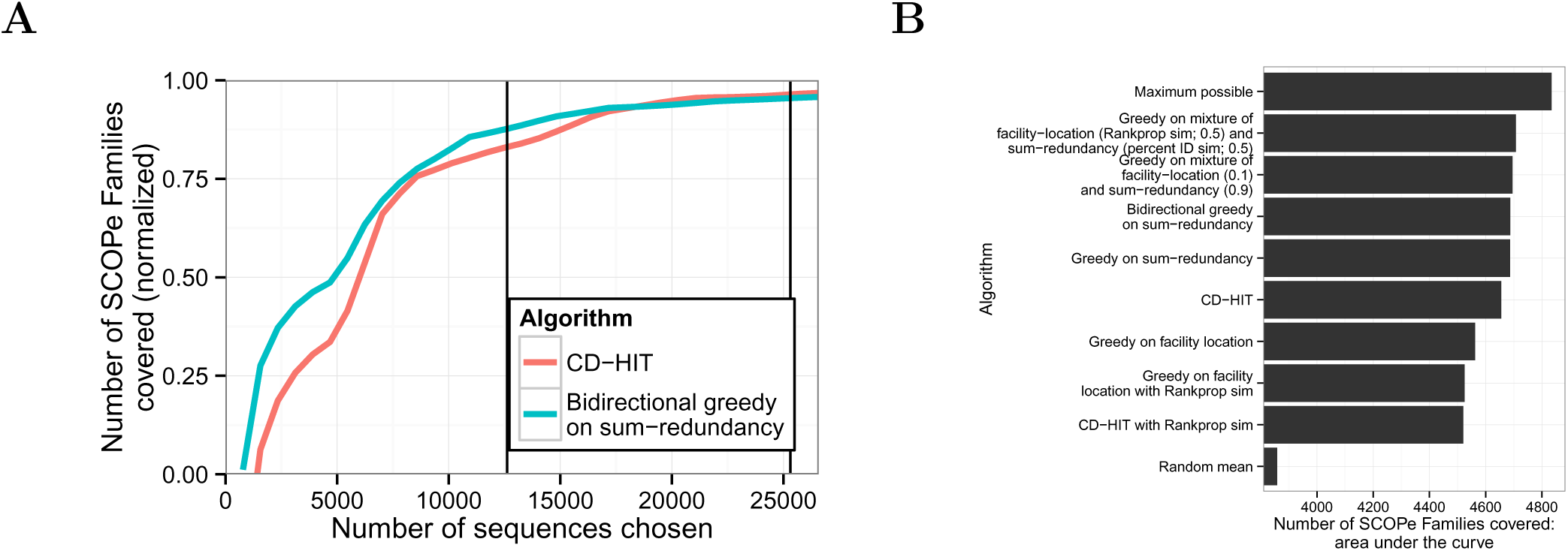
Coverage of SCOPe families. (A) The vertical axis indicates is the number of families with at least one representative, normalized to the range [0,1] using the formula 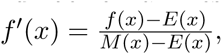 where *x* is a subset size, *f*(*x*) is the value for the method in question on that subset size, *E*(*x*) is expected (random mean) and *M*(*x*) is maximum possible (defined as *M*(*x*) = *x* if *x* < *F*, and *M*(*x*) = *F* otherwise, where *F* = 4,994 is the total number of SCOPe families). Vertical lines correspond to the sizes of subsets selected by CD-HIT using 40% and 90% sequence identity thresholds. (B) Area under the curve comparison. Bars indicate selection algorithms, ordered by their area under the curve. Horizontal axis indicates area under the “family” coverage curve, calculated as follows. Let *V* be the full set of sequences, *i* ∈ 1… |*V*| be a subset size, *m* be a particular method, 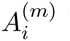 be the subset of size *i* chosen by *m*, and *f*(*A*) be the number of SCOPe Families covered by *A*. Area under the curve = 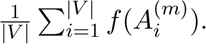 For subset sizes for which we did not compute a subset, *f*() is imputed with linear interpolation.

### 3.3 A mixture of submodular objective functions outperforms any single component objective

Although optimizing sum-redundancy performs best on the SCOPe-based evaluation metric for large subsets, we found that optimizing a different function resulted in better performance for very small subsets (<5,000 sequences; Figure 4A). Specifically, this good performance resulted from (1) optimizing a function called *facility-location* instead of sum-redundancy, and (2) using the similarity function used by the Rankprop method [Weston et al., 2004] that takes into account the statistical confidence of an alignment, instead of percent identity. The facility-location function is defined as f_facility-location_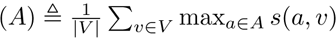 where *V* is the set of all sequences. Intuitively, this function takes a high value when every sequence in *V* has at least one similar representative in *A*. This function makes up for a weakness of redundancy-minimizing objectives such as sum-redundancy: such functions cannot differentiate representative sets with no detectable similarity between any representatives. In contrast, facility-location can differentiate sets of unrelated representatives by how well they cover unpicked sequences.

**Figure 4:**
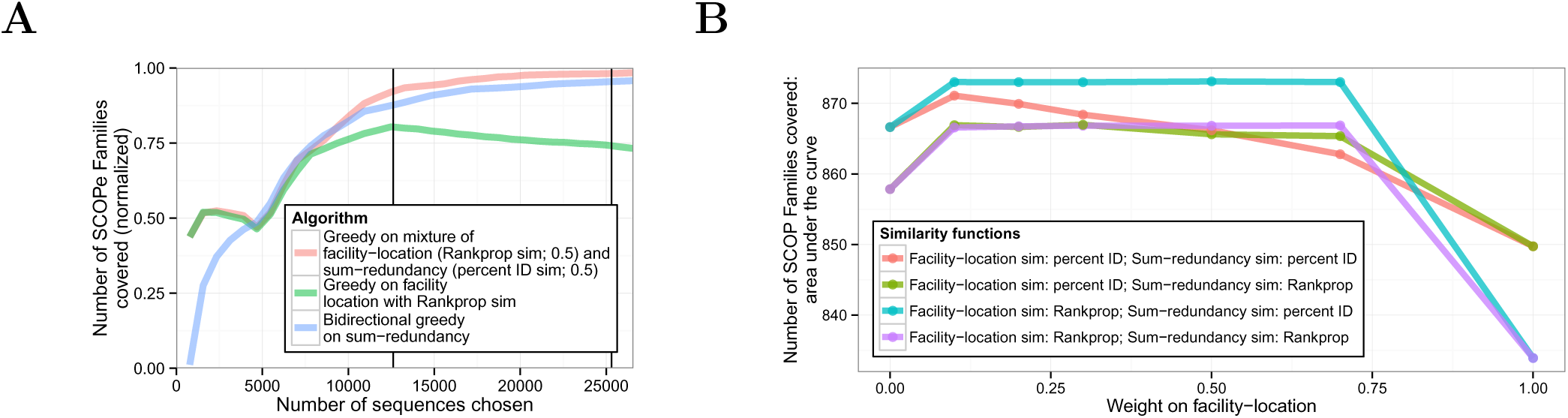
(A) Same as 3A, but including mixture objectives. (B) Comparison of mixture weights. Each mixture objective is a combination of facility-location and sum-redundancy objectives, each with either a percent ID or Rankprop similarity function. Horizontal axis indicates weight on facility-location (weight on sum-redundancy equals one minus the weight on facility-location). Line color indicates mixture objective. Vertical axis indicates area under the curve for SCOPe family coverage (see Supplementary Figure 5). To avoid overfitting, the statistics in (A) were computed on the subset of sequences with the SCOPe Class “All beta proteins”.

These observations suggest that sum-redundancy with the percent identical similarity metric and facility-location with the Rankprop similarity metric may provide different types of information. To test this hypothesis, we optimized a mixed objective function using the greedy algorithm and evaluated the resulting representative subsets against the SCOPe evaluation metric. We found that this mixture performed much better on SCOPe coverage than either function individually, both for small and large subsets (Figures 3B and 4A). Moreover, this performance is quite insensitive to mixture weight (Figure 4B). These results show that a mixture of objective functions can better represent the quantity of interest than a single function. Optimizing mixtures of objective functions this way is very natural using the submodular optimization framework, but would be very difficult in a non-optimization-based framework.

### 3.4 Representative subset selection methods perform better than re-purposed clustering methods

An alternative to applying a representative subset selection method is to apply a clustering method and then choose a representative from each identified cluster. Using the SCOPe evaluation metric, we compared representative subset selection using submodular optimization to eight variants of this clustering-based approach. We compared two similarity functions: percent identity and Rankprop; three clustering methods: *k*-medoids++, affinity propagation and agglomerative hierarchical clustering; and two methods for choosing a representative from each cluster: “random” and “central”. Central representative selection chooses the cluster member with the highest total similarity to the other cluster members (i.e. it optimizes a facility-location objective for a single item). Random representative selection chooses a random member. Note that *k*-medoids++ is identical to optimizing facility-location with the submodular greedy algorithm.

We found that none of these clustering methods performed as well as either the submodular optimization-based methods or the threshold heuristic (Figure 5 and Supplementary Figure 8). The exception, as noted above, is that optimizing facility-location with the Rankprop similarity metric performs well for small representative sets. This is likely because clustering methods aim to optimize the quality of the entire cluster, whereas representative set selection methods focus solely on identifying good representatives.

**Figure 5:**
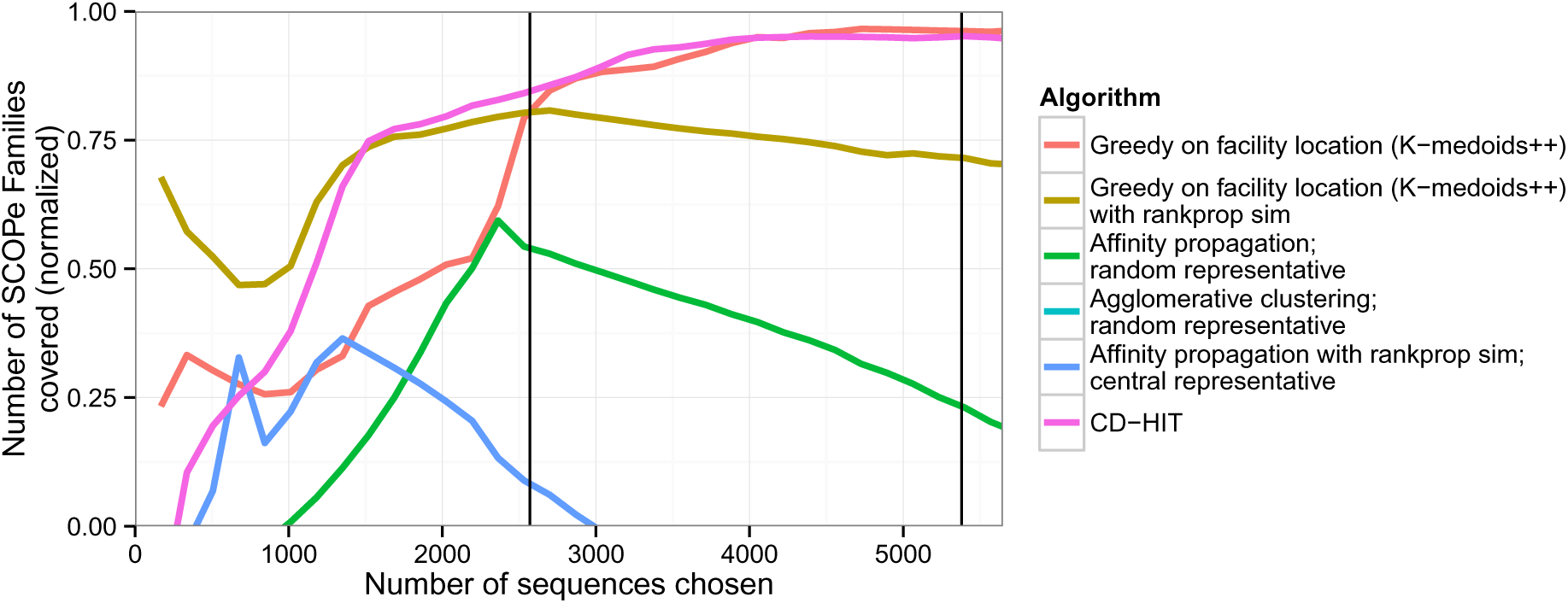
Performance of clustering methods repurposed for representative set selection. Plot is same as Figure 3, but includes clustering-based methods instead of submodular optimization-based. Results are computed on our development set, composed of the subset of the data (˜21%) with the SCOPe Class “All beta proteins”.

## 4 Discussion

The submodular optimization framework we present here results in theoretically guaranteed performance in some circumstances, performs well in practice, and has the flexibility to express a wide variety of desired properties of subsets of sequences. This approach is therefore preferable to existing, widely used, heuristic methods for selecting representative subsets of biological sequences. In this work we have explored only a small number of submodular functions; the submodular optimization framework supports many other objective functions that may provide even better results, including those based on cluster coverage, those based on the sequences directly rather than alignments, and others. Because the SCOPe evaluation function is itself submodular, a submodular objective function could also be efficiently learned from training data to correlate well with it. Moreover, this application may serve as a model for how submodular optimization can be applied to other discrete problems in biology, such as selecting representative sets of protein structures, genomic loci, or mass spectra.

## Acknowledgements

This work was supported by the National Institutes of Health award R01 CA180777.

